# MUSiCC: A marker genes based framework for metagenomic normalization and accurate profiling of gene abundances in the microbiome

**DOI:** 10.1101/009407

**Authors:** Ohad Manor, Elhanan Borenstein

**Author notes:** To whom correspondence should be addressed: Address: Foege Building S103B, Box 355065, Seattle WA 98195.

## Abstract

Functional metagenomic analyses commonly involve a normalization step, where measured levels of genes or pathways are converted into relative abundances. Here, we demonstrate that this normalization scheme introduces marked biases both across and within human microbiome samples and systematically identify various sample-and gene-specific properties that contribute to these biases. We introduce an alternative normalization paradigm, MUSiCC, which combines universal single-copy genes with machine learning methods to correct these biases and to obtain a more accurate and biologically meaningful measure of gene abundances. Finally, we demonstrate that MUSiCC significantly improves downstream discovery of functional shifts in the microbiome.

MUSiCC is available at http://elbo.gs.washington.edu/software.html.

## Introduction

The study of naturally occurring microbial communities through shotgun metagenomic assays has become a routine procedure in recent years [1–6]. Such assays are used, for example, to catalog the collection of genes in the metagenome, to estimate their abundances, and ultimately, to characterize the functional capacity of the community [1, 3, 7, 8]. This process involves two steps: First, genomic DNA is extracted from the sample and sequenced using next-generation technologies. Next, sequenced reads are aligned to a database of reference genes or genomes, and the number of reads that map to each gene is used as a proxy for its abundance in the sample [7, 9, 10]. Clearly, however, the resulting read counts are highly dependent on the sequencing depth in each sample, and some normalization method is required to allow comparison across samples. This is most commonly achieved by a simple *compositional normalization* process, whereby the obtained abundance value associated with each gene is divided by the sum of abundance values for all genes identified in the sample (e.g., [2, 11]). The resulting normalized value therefore represents a measure of *relative abundance* and is used in subsequent comparative analyses of the samples.

This normalization scheme, however, while extremely prevalent, has several fundamental weaknesses that may influence downstream analysis and ultimately impact the identification of functional shifts across samples. First, the resulting relative abundance values are unitless and do not necessarily represent a meaningful biological quantity. Second, in this normalization scheme, the scaled abundance of each gene crucially depends on the measured abundances of all other genes. As many different sample-specific factors could affect these quantities, abundance values could be disproportionately scaled in different samples, dramatically biasing any downstream comparative analysis. Compositional normalization is also associated with several statistical drawbacks and may give rise to misleading patterns [4, 12]. For example, as a marked increase in the abundance of one element decreases the apparent *relative* abundance of other invariant elements, this normalization scheme tends to induce spurious correlations between various elements in the sample. As a result, comparative analyses of these values across samples may be hard to interpret. These drawbacks call for an alternative normalization procedure, one that can produce accurate and easy to interpret abundance measures that can be reliably compared across samples.

Notably, a few previous metagenomics-based studies have already highlighted the challenges involved in compositional normalization. Specifically, studies of species composition have previously demonstrated that compositional normalization of taxonomic data could both mask true correlations between pairs of taxa and introduce false correlations [13–16]. Other studies of oceanic communities have further emphasized the biases introduced by compositional normalization of environmental metagenomic samples, specifically highlighting the potential contribution of the average genome size in each sample to these biases [17–19]. To date, however, the impact of compositional normalization on functional metagenomic studies of the human microbiome has never been shown or characterized, nor have the various sample-specific properties that may contribute to inaccuracies in abundance measures. Furthermore, previous studies of environmental metagenomes that aimed specifically to address genome-size induced bias still failed to provide biologically meaningful and interpretable measures of gene abundance.

Finally, even within each sample, various gene-specific properties may bias measured abundances. Compositional normalization, or for that matter, any normalization scheme that applies an identical processing protocol to all genes, will inevitably fail to account for such errors. Indeed, to date, no attempts to characterize or correct *within*-sample biases in genes’ abundances have been introduced, potentially neglecting important factors that may further contribute to inaccuracies in gene abundance measures.

In this study, we analyze samples from the Human Microbiome Project (HMP [2, 7, 11, 20]), as well as samples from two additional independent studies of the human gut microbiome [4, 11, 21], and demonstrate that compositional normalization has a clear and measurable effect on the obtained metagenomic functional profiles. We specifically show that this normalization protocol induces spurious inter-sample variation in the calculated abundances of various genes across samples from the same body site, across samples from different body sites, and across samples from different studies. We identify three sample-specific properties that play a key role in generating this spurious variation, including the average genome size, species richness, and mappability of genomes in the sample and suggest a simple and more biologically meaningful normalization method that aims to quantify the typical genomic copy number of each gene in the sample. We additionally show that gene-specific properties, such as sequence conservation and nucleotide content, further induce spurious intra-sample variation in the measured gene abundances, and provide an additional machine learning-based method to correct these inaccuracies. Finally, we demonstrate that our correction paradigm indeed provides improved abundance estimates and has clear and significant benefits for downstream comparative analyses. Specifically, we show that our method aids in the discovery of differentially abundant genes (and corrects false discovery of invariant genes), markedly increases the power to detect disease-associated pathways, and supports cross-study comparative analyses. Combined, these benefits allow us to move towards a more rigorous and unbiased estimate of the average genomic copy number of each gene in human microbiome samples, facilitating an accurate identification of functional shifts that may be associated with disease.

## Results and discussion

### Spurious inter-sample variation in HMP samples and its determinants

To examine the impact of compositional normalization on the analysis of human microbiome samples, we first identified a set of 76 *universal single-copy genes* (USiCGs; see Methods and Table S1). These genes are found in the genomes of almost all microorganisms, and generally at a single copy and therefore represent a proxy for invariant genomic elements. Ideally, therefore, since the content of each metagenome represents a simple linear combination of the genomic content of the constituent species’, one would expect such USiCGs to also be invariant across metagenomes. Put differently, comparative metagenomic analyses, which aim to identify shifts in the genic composition of the microbiome, could be considered accurate only if genes that are universally present in every member species in every sample exhibit no variation across samples. However, examining the relative abundances of USiCGs in metagenomic samples from the HMP (Methods), we found a marked variation both between and within body sites (Figure 1; Figure S1). For example, the median relative abundance of USiCGs in tongue dorsum samples is on average 1.3-fold higher than the median relative abundance of these genes in retroauricular crease samples. Similarly, the median relative abundance of USiCGs in some stool samples is 1.9-fold higher than in other stool samples. Importantly, this spurious variation in the observed abundance of USiCGs indicates that the calculated abundances of other genes may also be biased, potentially affecting any downstream comparative analysis and the correct identification of differentially abundant pathways.

**Figure 1.**
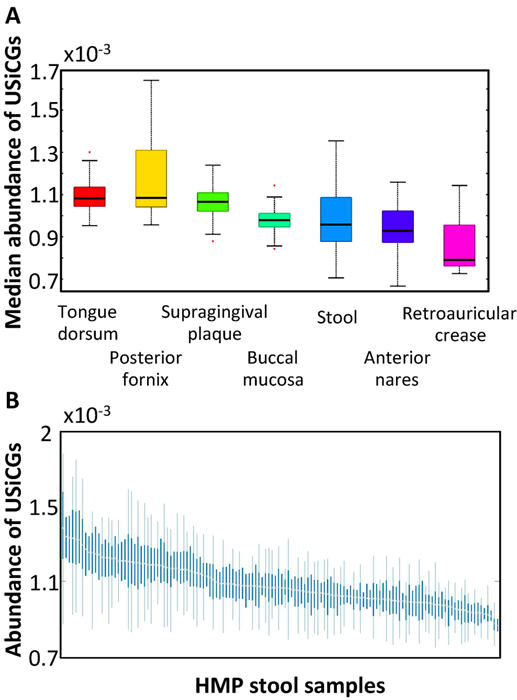
Spurious inter-sample variation across HMP metagenomic samples. **(A)** Variation in the relative abundance of USiCGs across different body sites (P<10^−300^, Kruskal-Wallis test). For each sample, the median relative abundance of the 76 USiCGs was used as a representative value. Boxes denote the interquartile range (IQR) between the first and third quartiles with the black line inside each box denoting the median. Whiskers extend to the lowest and highest values within 1.5 times IQR and outliers are marked in red. **(B)** The relative abundance of USiCGs across HMP stool samples. Each box illustrates the distribution of the relative abundances of the various USiCGs within each sample. Box range, whiskers, and outliers are defined as in panel A. See also Figure S1 for the distribution within all other body sites.

Notably, the variation in the abundance of the various USiCGs appears to be consistent and sample-specific; in certain samples the abundances of all USiCGs tend to be high whereas in others the abundances of all USiCGs tend to be low (Figure 1; Figure S1). We therefore set out to identify sample-specific properties that may contribute to this spurious variation in USiCGs’ abundances. One obvious candidate is the average genome size in the sample. Indeed, previous studies have demonstrated that the average size of genomes vary substantially across samples [22, 23]. Since USiCGs, by definition, appear in one copy in each genome, as the average genome size in the sample increases, the *fraction* of the sample’s DNA that originates from USiCGs becomes smaller, and accordingly, the calculated relative abundance of USiCGs decreases. This further highlights the problematic nature of compositional normalization, wherein an increase in the abundance of some element in the metagenome brings about an artificial and biologically meaningless decrease in the calculated relative abundances of *all* other elements in the sample. Previous studies of marine microbiome samples have indeed suggested that the average genome size can account, at least partially, for bias in the relative abundance of a small set of invariant genes across samples [13, 15, 18]. To examine whether such association between spurious variation and the average genome size can also be observed in the human microbiome, we used a recently introduced computational method [17, 24] to estimate the genome size of each operational taxonomic unit (OTU) in every sample (Methods). We then calculated the average genome size in each sample, weighting the genome size of each OTU by the OTU’s relative abundance in the sample. Comparing these average genome sizes with the median relative abundance of USiCGs across all body sites, we indeed found a strong negative correlation (R=-0.65, P<10^−21^, Pearson correlation test), with the relative abundance of USiCGs tend to be lower in samples with larger genomes (Figure 2A). A similar correlation is clearly observed also across samples from the same body site (e.g., R=-0.82, P<10^−15^, stool; R=-0.6, P<10^−3^, tongue dorsum; Pearson correlation test, Figures 2B and S3B).

**Figure 2.**
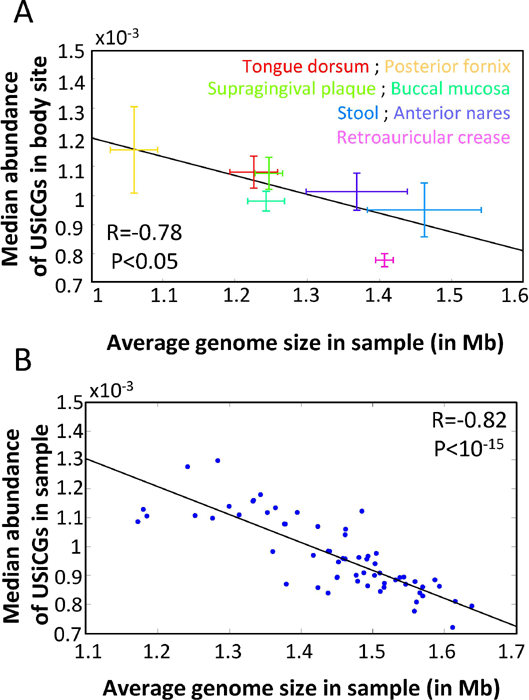
The median abundance of USiCGs across HMP samples is negatively correlated with the average genome size both **(A)** across different body sites (R=-0.78, P<0.05), and **(B)** across stool samples (R=-0.82, P<10^−15^). See Methods for more details on average genome size estimation. In panel A, each cross represents the median and the median absolute deviation across all samples from each body site. In panel B, each point represents a single stool sample. Regression lines are illustrated in black.

We next set out to examine additional, yet to be identified sample-specific properties that may further contribute to biasing observed relative abundances. We specifically focused on two such candidate properties: the species richness of the sample, and the mappability of species in the sample to fully sequenced reference genomes. Both these properties may influence the fraction of reads that map to known genes (or functions) and could consequently impact the relative abundance of reads that map to, for example, USiCGs. Specifically, as species richness increases and the community includes rarer and more poorly characterized species, and as the overall mappability of community members to fully sequenced species decreases, more and more genes in the genomes of the various community members may not have a close ortholog in the database and reads originating from these genes may fail to map to any known orthology group. In contrast, reads that originate from USiCGs (that are by definition present in every genome and are well conserved) or from other, relatively common genes, will likely still map successfully to their known orthologs, and their *relative* abundance will accordingly appear higher. Using the number of OTUs identified in a sample as a measure for species richness, and the average evolutionary similarity between each OTU in a sample to its nearest sequenced reference genome as a measure of mappability (Methods), we indeed found that both these properties correlate with the relative abundances of USiCGs. Specifically, in the gut microbiome, for which the largest number of HMP samples was available, the correlation between the median relative abundance of USiCGs and species richness was 0.44 (P<10^−3^; Pearson correlation test) and the correlation between USiCGs’ abundance and mappability was -0.85 (P<10^−17^; Pearson correlation test). Evidently, as the richness of species in the sample decreases or their similarly to other sequenced reference genomes increases, the measured relative abundance of USiCGs increase as well (Figure S2A-B). Importantly, although these three sample-specific properties (namely, average genome size, species richness, and mappability) are inter-correlated, each property is significantly correlated with the abundance of USiCGs even when controlling for the other two properties (see Methods), suggesting that each such property has an independent impact on USiCGs variation and is not just a correlated byproduct (Table 1; Figure S2C-D). A similar pattern was also observed in the tongue dorsum microbiome (Figure S3; other sites did not have enough samples available for such an analysis)

**Table 1A.** Correlations and controlled correlations between the abundance of USiCGs and various sample-specific properties in the stool samples.

| Sample-specific property | Correlation with USiCGs abundance | Controlled correlation with USiCGs abundance* |
| --- | --- | --- |
| Genome size | -0.82 ($P < 10^{-14}$) | -0.46 ($P < 10^{-3}$) |
| Mappability | -0.85 ($P < 10^{-17}$) | -0.58 ($P < 10^{-6}$) |
| Species richness | 0.44 ($P < 10^{-3}$) | 0.37 ($P < 0.003$) |

\* Partial correlation, controlling for the other two properties (see Methods)

**Table 1B.** Correlations between sample-specific properties.

| Property 1 | Property 2 | Correlation |
| --- | --- | --- |
| Genome size | Mappability | 0.8 ($P < 10^{-300}$) |
| Genome size | Species richness | -0.25 ($P < 0.04$) |
| Mappability | Species richness | -0.27 ($P < 0.02$) |

### Spurious inter-sample variation across different studies

Having demonstrated clear spurious variation in the abundance of USiCGs across HMP samples, we next examined whether such variation can also be observed in the abundance of these genes across independent studies. We therefore obtained metagenomic data describing the human gut microbiome of healthy individuals from 3 different studies, including 134 samples from the Human Microbiome Project (*HMP*) [2, 19], 60 samples from a study of the gut microbiome in inflammatory bowel disease in European individuals (*IBD*) [4, 25], and 174 samples from a study of type 2 diabetes in Chinese individuals (*T2D*) [11] (Methods). We computed the median abundance of USiCGs in each sample and compared these values across studies. We again found marked variation in the relative abundance of USiCGs, with up to a 3-fold difference between studies (Figure 3A) and up to a 2.3-fold difference within a single study (Figure 3B).

**Figure 3.**
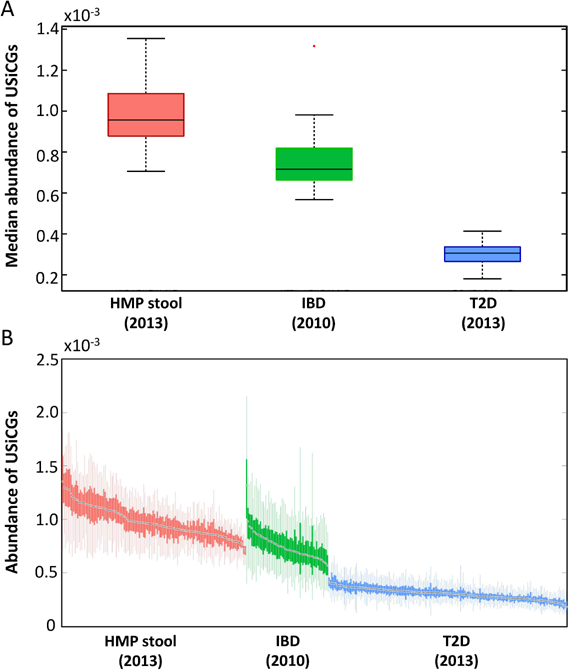
Spurious inter-sample variation in the functional metagenome of healthy gut microbiome samples across three independent studies. The median relative abundance of USiCGs across the different studies **(A)** and the relative abundance of USiCGs within each study **(B)** are illustrated. Boxes and whiskers are defined as in Figure 1.

We further wished to examine whether the same sample-specific properties that we identified above as contributing to this bias in HMP samples play a role also in the observed spurious variation across the different studies. However, as full taxonomic data (in the form of OTU counts) was only available for HMP samples, a direct comparison was not feasible. Instead, we therefore searched for a sample-specific property that was available for all 3 studies and that could serve as a statistical proxy for the OTU-based properties examined above. Using the HMP dataset, we found that the total number of assembled contigs in a sample was highly correlated with the average genomic size, species richness, and average mappability in the sample (Pearson correlation R=-0.52, R=0.55, and R=-0.53, respectively). Since all 3 studies performed de-novo contig assembly, this measure could be used in lieu of the more explicit sample-specific properties utilized above. Comparing the number of contigs per sample in the various studies with the median relative abundance of USiCGs, both between and within the different studies, we indeed found a similar trend as observed above. Specifically, the study with the highest number of assembled contigs (HMP; ~119,000 contigs on average per sample) was also the one with the highest median relative abundance of USiCGs (8.65 × 10^−4^), whereas the study with the fewest contigs (T2D; ~27,000 contigs on average per samples) was the one with the lowest median relative abundance of USiCGs (2.63 × 10^−4^). Similarly, a modest to strong correlation was found between the number of assembled contigs and USiCGs’ abundance across the samples within each study (R=0.6, R=0.11 and R=0.28 for HMP [2], IBD [4], and T2D [11], respectively).

These findings suggest that spurious variation in the relative abundance of USiCGs is also evident across multiple independent studies and may be accounted for by similar sample-specific properties. Importantly, this implies that compositional normalization may hinder future attempts to pool datasets from multiple studies in order to increase the power to discover functional variation in the human microbiome (and see also our cross-study comparative analysis below).

### Inter-MUSiCC: Correcting spurious inter-sample variation

To correct the spurious inter-sample variation demonstrated above, we present here a fundamentally different normalization scheme. Clearly, with the identification of sample-specific properties that contribute to spurious variation, one potential approach for addressing this challenge would be to carefully characterize how each sample-specific property affects normalized abundances and apply some mathematical formulation to reverse this effect. Previous attempts to specifically correct the impact of average genome length indeed took this approach [18, 19]. However, considering the multiple sample-specific properties identified in the previous section and their complex intertwined contribution to spurious variation, we took a more direct approach that corrects spurious sample-specific variation, regardless of its determinants, and provides a biologically intuitive and meaningful measure for gene abundances in a metagenome (and see also our analysis of simulated samples below). Specifically, our method aims to describe the abundance of each gene in the microbiome, not as its relative abundance in the sample, but rather as the average copy number of this gene (with respect to some copy number proxy as described below) across all microbial cells in the sampled community. The functional profile obtained by this normalization method for a given sample can accordingly be conceived as a description of the gene content of a ‘typical microbe’ from that sample. This profile therefore has a clear biological interpretation, and can be reliably and meaningfully compared across samples.

In practice, this normalization paradigm is achieved by using the same set of USiCGs described above as the yardstick for estimating the average copy number of all other genes. Technically, the measured abundance of each gene in a sample (after correcting for gene length) is divided not by the sum of all abundances, but rather by the median abundance of the USiCGs in the sample. With this correction, USiCGs in every sample have a consistent median value of one (as expected by their definition), and all other genes are measured in comparison to these USiCGs. We therefore term this normalization method *Inter-sample Metagenomic Universal Single-Copy Correction (Inter-MUSiCC).*

### Evaluating the impact of Inter-MUSiCC on functional metagenomic profiles

Inter-MUSiCC, by definition, corrects the spurious inter-sample variation we demonstrated above in USiCGs. For this normalization scheme to be useful, however, it should clearly also yield improved abundance estimates for all other genes. We focus on HMP stool samples, owing to the large number of samples available and the ability to examine also the impact of MUSiCC on multiple studies of the gut microbiome. Focusing on a single body site also allows us to assess the performance of MUSiCC on a more homogenous set of samples and therefore to evaluate MUSiCC’s ability to correct even the relatively fine variation observed in such samples. Notably, however, as the actual average copy number of each gene in these samples is unknown, evaluating the impact of inter-MUSiCC and assessing whether it produces reasonable estimates of average copy numbers in these metagenomes is a challenging task.

To address this challenge, we considered the set of genes that occur in only one OTU per sample (Methods). For such OTU*-Specific Genes* (OSGs), the relationship between the abundance of each gene and the abundance of the corresponding OTU is not complicated by the presence of other OTUs in the sample, making it easier to evaluate the impact of Inter-MUSiCC on the corrected abundance values. Specifically, as each OSG occurs in only one OTU, clearly, the abundance of the OSG across the various samples should positively correlate with the abundance of the respective OTU. If Inter-MUSiCC indeed provides a more accurate estimation of gene abundance compared to compositional normalization, this correlation between OSGs’ and OTUs’ abundances should improve with the application of Inter-MUSiCC. Since many OTUs observed in each sample are not yet associated with a fully sequenced genome, we used a recently introduced tool to predict the genomic content of each OTU (Methods; [24]). In total, we identified 3,821 OSGs in 993 OTUs across 65 HMP stool samples. For each such OSG, we calculated the correlation between its abundance across the various samples and the abundance of its respective OTU with and without the application of Inter-MUSiCC. We found that Inter-MUSiCC indeed significantly increased the average correlation between the abundances of OSGs and their respective OTUs (P<10^−32^, paired t-test; Methods), suggesting that the abundance values obtained by Inter-MUSiCC more accurately reflect gene abundances. To further confirm the beneficial impact of Inter-MUSiCC on the abundance of OSGs, we additionally examined the ratio of OSGs’ (and OTUs’) abundances across pairs of samples. Such a ratio-based analysis may cancel out various factors that confound the relationship between OSGs’ and OTUs’ abundances (such as false detection of OSGs), further highlighting the accuracy of calculated OSGs’ abundances. Specifically, as each OSG occurs in only one OTU, the ratio between its abundance in one sample to its abundance in another should be similar to the ratio between the abundances of the respective OTU in these two samples. For example, if a certain OTU exhibits a 3-fold increase in abundance between two samples, any OSG that is associated with only this OTU in these two samples is also expected to exhibit a 3-fold increase in abundance. To confirm this, for each case where the same OSG appeared in two different samples, we quantified the difference between the fold-change in the OSG abundance between the two samples and the fold-change in the OTU abundance between these samples (Methods). We found that both when using compositional normalization and when using Inter-MUSiCC, OSGs exhibit on average a higher fold-change than expected from the fold-change in their respective OTU abundances. This may reflect subsampling issues, wherein low-abundance OTUs can be accurately measured through 16S sequencing, while the abundance of their genes may still be underestimated by shotgun metagenomic sequencing. Yet, with Inter-MUSiCC, OSGs’ abundance fold-change was significantly closer to the measured fold-change in OTU abundance (P<10^−315^, paired t-test), suggesting that at least some of the bias in the abundances of each gene was corrected.

Finally, since compositional normalization is known specifically to introduce false correlations between normalized elements [12], we set out to examine the impact of Inter-MUSiCC on pairwise gene correlation. As before, a gold standard for the true correlation structure in the human microbiome is not available. However, since metagenomes represent a large collection of genomes, one could expect that pairs of genes whose occurrences across genomes are highly correlated (*i.e.,* they tend to either both be present in a genome or both be absent), would also be correlated in their abundance across metagenomes. We therefore compared for each gene the correlation of its occurrence and the occurrence of every other gene across fully sequenced genomes in the KEGG database [1, 3], with the correlation of it abundance and the abundance of every other gene in HMP stool metagenomic samples (Methods). We found that with Inter-MUSiCC the similarity between the genomic and metagenomic correlation structure is significantly higher than with compositional normalization (P<10^−121^, Student’s t-test), again suggesting that Inter-MUSiCC better captures the abundance levels of the various genes in the metagenome.

### Spurious intra-sample variation and its determinants

In the previous sections, we demonstrated that several sample-specific properties could bias the measured relative abundances of the various genes in each metagenomic sample, giving rise to spurious inter-sample variation. However, examining the relative abundance of USiCGs within a single sample (after normalizing for their length), we also find marked variation *within* samples. Importantly, this intra-sample variation is highly consistent across samples, with some USiCGs regularly exhibiting higher abundance than others (Figure 4). Since every genome in the sample encodes the same number of copies of each of these genes (namely, a single copy), this intra-sample variation clearly does not represent true variation in the composition of the sample. Moreover, the consistency in this variation suggests that it can be attributed to gene-specific properties that systemically bias the measured abundance.

**Figure 4.**
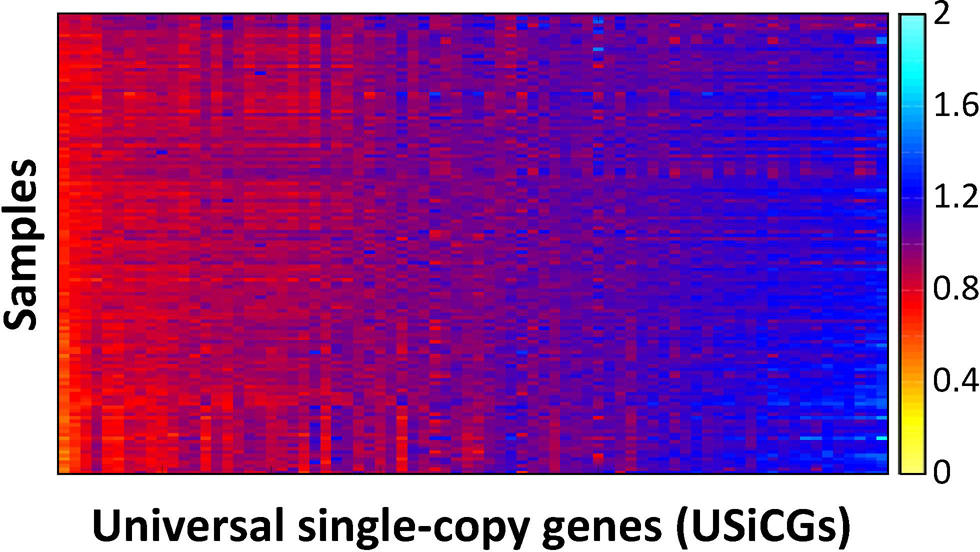
Spurious intra-sample variation in the abundance of USiCGs is consistent across samples. The heatmap illustrates the abundance of each of the 76 USiCGs (x-axis) across 134 HMP stool samples (y-axis). The color gradient denotes the abundance after correcting for spurious inter-sample variation (using Inter-MUSiCC), and hence represents the average copy number of the USiCG in the sample.

To test this hypothesis, we collected a set of 35 gene-specific properties, including various conservation features (across the gene orthology group) and nucleotide content measures (Table S2; Methods). Focusing again on HMP stool samples, we identified a subset of these properties that were significantly correlated with intra-sample variation in USiCGs (measured as the fold-change between the abundance of each USiCG and the average abundance of all USiCGs in the sample; Table S2). For example, USiCGs with high GC content tended to exhibit lower abundance compared to USiCGs with low GC content. Similarly, USiCGs with highly variable length (measured as the standard deviation in the length of the gene across all members of the gene orthology group) were more likely to have lower abundance compared to USiCGs with a more consistent length. To examine the predictive power of such gene-specific properties on spurious intra-sample variation and to account for the potentially strong dependencies between the various properties, we further used a machine learning approach with L_1_ regularization to obtain for each sample a linear model that links these gene-specific properties to observed variation (Methods). Remarkably, we found that this linear model correctly predicts and can correct on average >50% of the observed intra-sample variability in USiCGs’ abundance on held-out test data across HMP stool samples (Figure 5A). Moreover, examining which gene-specific properties were selected by the linear model in each sample, we found a clear agreement between models across the various samples (Figure 5B), highlighting the robustness of this model. Among the properties that were repeatedly selected by the model were the median GC nucleotide content of the gene orthology group and the number of species in which this gene could be detected using KEGG’s hidden-Markov-model.

**Figure 5.**
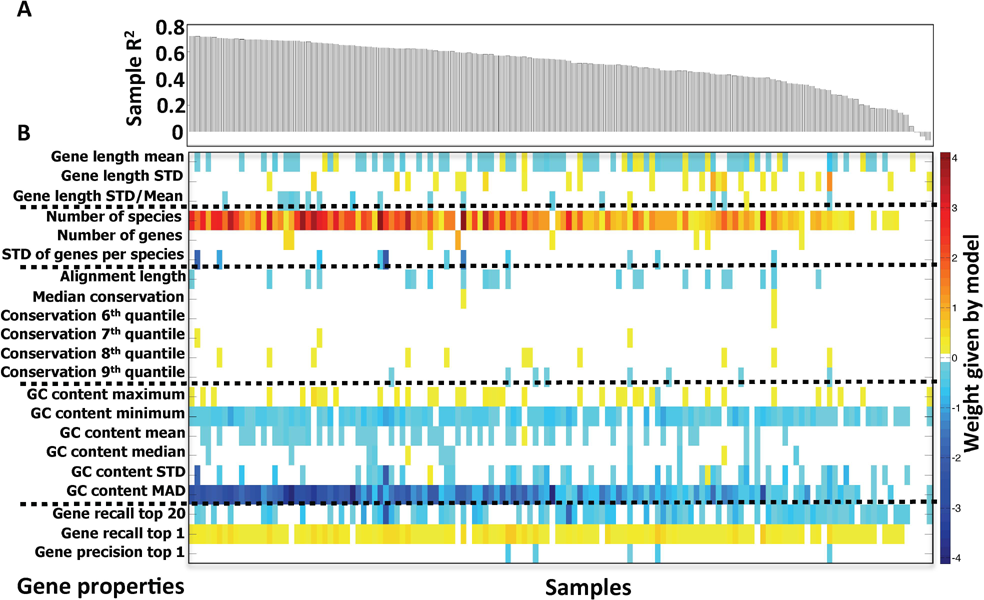
L_1_ regularized linear modeling of spurious intra-sample variation in the abundance of USiCGs. **(A)** The proportion of explained variation (R^2^) in each HMP stool sample on held-out test data. **(B)** The weights (color gradient) assigned to the various gene-specific properties by an L_1_-regularized linear regression model across the different samples.

### Intra-MUSiCC: Correcting spurious intra-sample variation and evaluating its impact on functional metagenomic profiles

Our findings above suggest that spurious intra-sample variation in USiCGs could be corrected by learning a predictive linear model that accounts for the impact of various gene-specific properties on measured gene abundances. Assuming that such spurious intra-sample variation also occurs in other, non-USiCG genes, we can use the USiCGs-based model as a proxy for the effect of gene-specific properties and apply this model to correct the measured abundances of all other genes in the sample. Since this method again relies on the set of USiCGs identified above, we term it *Intra-sample Metagenomic Universal Single-Copy Correction* (*Intra-MUSiCC*). Notably, this predictive model can be learned once (e.g., using a large set of metagenomic samples) and applied to every new sample without modification. In what follows, however, we used a somewhat more sophisticated approach wherein a predictive model is learned in each sample separately (based on the abundances of USiCGs in that sample), cross-validated on unseen USiCGs abundances, and used to correct variation within that sample. This approach assumes that the impact of gene-specific properties on measured abundances may slightly vary from sample to sample and aims to capture the potentially unique effect of gene-specific properties in the sample. We confirmed however that the more generic approach, in which the predictive model is learned once and applied to all samples, produced qualitatively the same results as reported below.

In the previous section, we already demonstrated that Intra-MUSiCC corrects much of the spurious intra-sample variation in USiCGs’ abundances (see Figure 5). However, to confirm that Intra-MUSiCC is a useful and applicable correction scheme, we next set out to validate that it indeed yields improved abundance estimates also for non-USiCGs genes, focusing, as before, on the set of HMP stool samples (see above). As was the case for our analysis of Inter-MUSiCC, this task is challenging since the true abundance values of non-USiCGs across samples is not available. We therefore used several different approaches, each focusing on a specific set of genes, to evaluate the performance of Intra-MUSiCC and to confirm that the corrected gene abundance values are indeed more accurate and more informative than the widely-used relative abundance values.

First, we identified a set of 72 genes that did not meet our criteria for USiCGs, but that could still be considered universal single-copy genes under a more relaxed set of requirements (Table S3; Methods). Specifically, such “semi”-USiCGs tend to have a single copy in the vast majority of genomes, but are not as prevalent as the USiCGs described above. While this set is expected to be more variable than USiCGs (due to true variation in their occurrence across genomes), we hypothesized that at least some of the observed intra-sample variation among these genes might be spurious and accounted for by the various gene-specific properties detected above. Indeed, we found that when using the USiCGs-based model (*i.e.,* Intra-MUSiCC) to correct the abundance values of these genes across all HMP stool samples, 17% of the variation in these semi-USiCGs was corrected. Notably, this reduction in variation is observed even though these semi-USiCGs were never used in the construction of the model.

Next, we focused on pairs of genes whose occurrence is highly correlated across genomes. Specifically, examining the gene content of all reference genomes in KEGG [1, 3], we identified 1074 pairs of genes whose presence/absence profiles agree across >95% of the genomes in which they appear (Methods). Importantly, we excluded all USiCGs (as well as the semi-USiCGs described above) from this analysis. For each such pair, we computed the average absolute difference in abundance across HMP stool samples, before and after applying Intra-MUSiCC. Clearly, high cooccurrence across genomes does not necessarily imply perfectly correlated abundance across metagenomes. A metagenome may, for example, include a relatively large proportion of exactly those genomes in which the two genes do not co-occur. Yet, it is reasonable to assume that such genomically co-occurring genes will tend to exhibit similar abundances across metagenomes. Accordingly, we hypothesized that the observed differences between the abundances of such gene pairs can be partly accounted for by spurious intra-sample variation that arises from gene-specific bias as opposed to true biological variation. In line with this hypothesis, we indeed found that both the mean and variance of the pairwise differences in the abundance of each of the 1074 gene-pairs were significantly lower with the application of Intra-MUSiCC (P<10^−66^ and P<10^−46^, for mean and variance, respectively; paired t-test for mean, paired F-test for variance).

Finally, we extended the analysis of genomically co-occurring genes, focusing on clusters (rather than pairs) of genes whose occurrence is highly correlated across genomes. We again used the set of KEGG reference genomes to identify 25 gene clusters, each consisting of at least 5 genes (with a total of 216 genes), whose pairwise presence/absence profiles agree in >95% of genomes (Table S4; Methods). Again, we assumed that such genomically co-occurring genes are likely to exhibit similar abundances across HMP gut metagenomic samples. We used Intra-MUSiCC to correct the abundance of each gene in the various clusters and compared the corrected abundance measures to the original relative abundance values in each cluster. Importantly, using clusters of genes rather than independent pairs allowed us to examine not only the pairwise difference in abundance but also the proportion of variance explained within each cluster. We found that on average Intra-MUSiCC corrected 8.2% of the variance within each gene cluster (Table S4). As an example, one such gene cluster contained all 8 genes in the F-type ATPase structural module. Although these 8 genes are highly consistent in their presence/absence patterns across reference genomes (median pairwise Jaccard similarity of ~99%), their abundances across HMP stool samples vary by up to ~2.1-fold on average (and >13.4-fold in some samples). Intra-MUSiCC (which notably employs a model learned on USiCGs alone and for which these 8 genes can be conceived as unseen data) corrected on average 32% of this variance.

### MUSiCC markedly enhances the discovery of functional shifts in the microbiome

Above, we demonstrated that MUSiCC successfully corrects both inter-and intra-sample spurious variation in gene abundances. Clearly, however, the key goal of any metagenomic normalization scheme is to facilitate comparative analysis and to enable the discovery of functional shifts in the metagenome that may be associated with a given host phenotype. Next, we therefore set out to examine whether MUSiCC (*i.e.,* the combined application of Inter-MUSiCC and Intra-MUSiCC) has a concrete and significant impact on such comparative analyses and whether it affects the identified set of differentially abundant genes or pathways in various settings.

We first examined the impact of MUSiCC on the detection of differentially abundant genes. Specifically, we used standard comparative analysis (Methods) to identify genes that exhibit differential abundance between HMP stool samples and tongue dorsum samples, with and without the application of MUSiCC to correct the abundances of genes. Comparing the obtained sets of differentially abundant genes, we found an overall agreement, with 90% of the detected genes identified both with and without MUSiCC. Notably, however, there were also substantial differences between the two sets, with 382 genes found to be differentially abundant only when using MUSiCC and 343 genes found to be differentially abundant only when using standard compositional normalization (Figure 6A). Interestingly, among the 382 genes identified only with MUSiCC, genes involved in Lipopolysaccharide biosynthesis – a known gut metabolic pathway [9, 10] – were over-represented (P<10^−3^, hypergeometric test). Conversely, the 343 genes detected only by compositional normalization showed over-representation of ribosomal genes (P<10^−8^, hypergeometric test) and in fact included many (36) of the 76 USiCGs. This clearly and effectively highlights the problematic nature of compositional normalization, wherein fundamentally invariant genes can be detected as differentially abundant due to the biases introduced by this normalization scheme. We observed a similar pattern also when comparing HMP stool samples to other HMP body sites, with an average of ~392 genes identified as differentially abundant in stool only when using MUSiCC (Figure S4).

**Figure 6.**
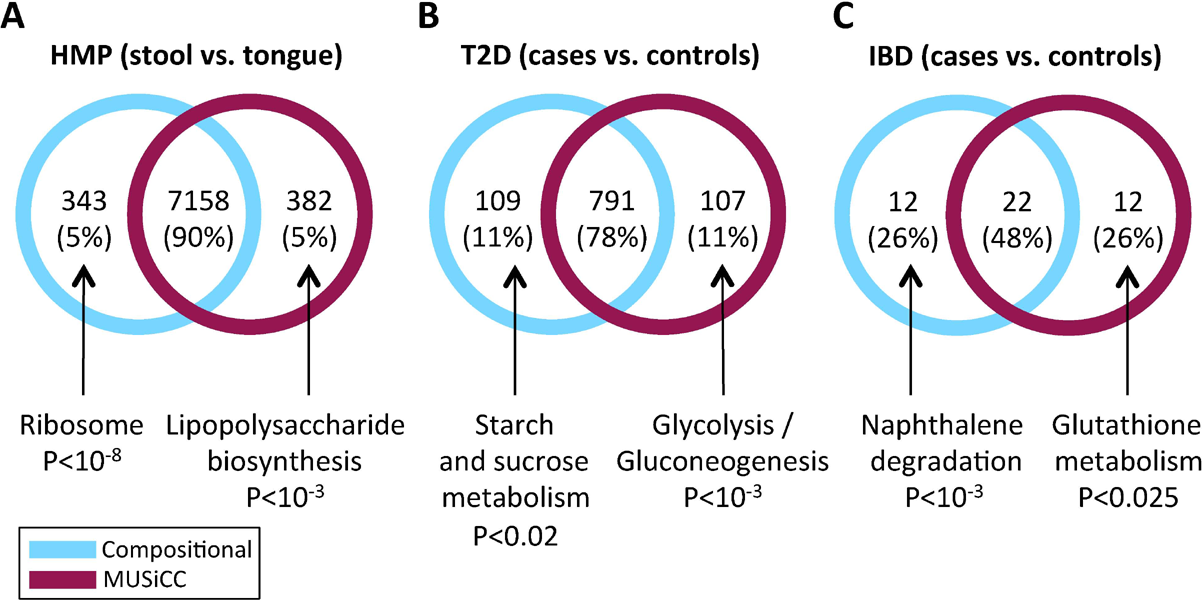
The impact of MUSiCC on the discovery of differentially abundant genes. The number of genes identified as differentially abundant between **(A)** HMP stool samples and tongue dorsum samples, **(B)** type 2 diabetes (T2D) cases and controls, and **(C)** inflammatory bowel disease (IBD) cases and controls, are illustrated. Each Venn diagram describes the overlap between the set of genes identified when using standard compositional normalization (cyan, left) and the set of genes identified when using MUSiCC (maroon, right). Pathways that are over-represented in the set of genes identified by only one of the two methods are listed.

Crucially, MUSiCC’s impact on the detected set of differentially abundant genes is observed not only when comparing different body sites in healthy individuals, but also when comparing disease cases vs. healthy controls. Specifically, analyzing the two disease-related studies discussed above (namely, the Type 2 diabetes study [11]; *T2D,* and the inflammatory bowel disease study [4]; *IBD),* a similar pattern was found, with many genes detected as differentially abundant by only one of the two normalization methods. In fact, in these datasets, the set of differentially abundant genes discovered with and without MUSiCC exhibit an even lower agreement, with only 78% and 48% overlap between the methods for T2D and IBD, respectively (Figure 6B-C). As before, the sets of disease associated differentially abundant genes discovered only by MUSiCC were enriched for pathways previously reported to be linked to T2D and IBD, including Glycolysis/Gluconeogenesis [13, 15] and Glutathione metabolism [17], respectively. These findings suggest that MUSiCC not only corrects various biases in metagenomic functional data, but can also lead to significant changes in the identification of disease-associated genes, pointing to novel candidates for microbiome-based intervention (and avoiding false discovery of other, clearly unrelated genes).

Moving beyond individual genes, we next set out to examine whether MUSiCC affects the discovery of pathway-level functional shifts in the microbiome. Such pathway-level analyses and the study of pathways identified as associated with a disease state is one of the most common downstream comparative analysis approaches (e.g. [7, 11, 20]). Following this paradigm, we conducted a comparative analysis of pathway abundances in the two disease-related datasets described above, to identify T2D-and IBD-associated pathways (Methods). Comparing the set of T2D-associated pathways identified when using standard compositional normalization to the set identified after applying MUSiCC, a clear impact was observed both on the number of pathways discovered and on their statistical significance (Figure 7A). Specifically, all 19 T2D-associated pathways identified with compositional normalization were also identified with MUSiCC, but in 13 of these 19 pathways (68%), the significance level was increased when using MUSiCC. Moreover, with MUSiCC, 17 additional T2D-associated pathways (that did not pass the significance threshold when using compositional normalization) were identified, including pathways previously reported as linked to T2D (e.g., methane metabolism [21] and Glycolysis [13, 15]). A similar pattern was observed in our analysis of the IBD dataset (Figure 7B). Specifically, all 7 pathways identified as IBD-associated with compositional normalization were also identified with MUSiCC, with 6 out of these 7 pathway (86%) becoming more significant with the application of MUSiCC. Similarly, 28 additional pathways were identified only with the application of MUSiCC, including previously reported IBD-associated pathways (such as Riboflavin metabolism [17]).

**Figure 7.**
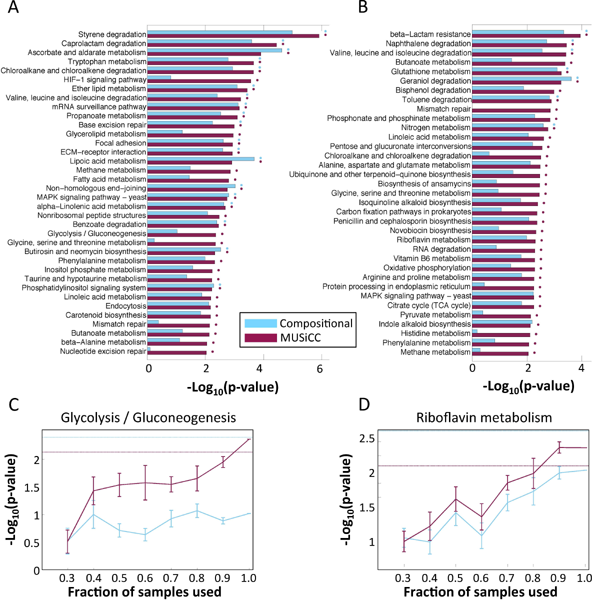
The impact of MUSiCC on the discovery of disease-associated pathways. Pathways identified to be associated with **(A)** type 2 diabetes, or with **(B)** inflammatory bowel disease, when using standard compositional normalization (cyan) or MUSiCC (maroon). Bars denote the significance level of the association. The dots to the right of each bar indicate whether this association reached significance with FDR<0.05 with compositional normalization (cyan) or MUSiCC (maroon). (**C-D**) Trend plots demonstrating the significance level of the association of the Glycolysis pathway with T2D (C) and of the Riboflavin metabolism pathway with IBD (D), as a function of the fraction of the samples used. Mean and standard errors using compositional normalization or MUSiCC are illustrated.

Considering the overall increase in the significance level of disease-associated pathways observed above with MUSiCC (Figure 7A-B) and since many disease-related studies can obtain or process only a limited number of samples, we further examined whether MUSiCC can indeed enhance the power to detect disease-associated pathways when data is limited. We therefore repeated the comparative analysis above, focusing on two disease-associated pathways of interest, and measured how significant the obtained association signal was when using only a subset of the available samples. Indeed, we found that with MUSiCC, significance is reached with far fewer samples than with compositional normalization (Figure 7C-D), suggesting that MUSiCC can successfully uncover novel disease-associated pathways that may be masked by noise or by sparse sampling when using standard compositional normalization. Using this disease-associated pathway discovery as the ultimate benchmark for the applicability of the various normalization methods, we additionally tested an alternative approach that was previously used to process a set of oceanic samples ([19]; Methods). We found that this method outperformed the standard compositional normalization approach but was nonetheless far less successful than MUSiCC and still failed to identify many of the relevant pathways as significant (Figure S5).

Finally, having shown that MUSiCC promotes the discovery of disease-associated pathways (and with fewer samples), we set out to examine whether it can also allow researchers to combine data from multiple independent studies. Pooling sample sets across studies could dramatically increase the amount of data available for future comparative analyses and has the potential to significantly enhance efforts to discover disease-associated shifts in the microbiome. Specifically, this approach could be used to harness a large collection of healthy samples (such as those obtained by HMP) as controls for many smaller disease-focused studies. Notably, however, we already identified above marked spurious variation between different studies of the human gut microbiome (see Figure 3), suggesting that a naïve pooling of data from multiple studies without careful correction of sample-specific biases may be challenging. Indeed, comparing the pathway-level functional profiles of the healthy samples from the three studies and performing a principal coordinate analysis (PCoA) to examine the variation in this pooled set of samples, it was clear that the vast majority of the variation is study specific (Figure 8A). In such settings, using healthy samples from one study as the set of controls for another study would potentially mask genuine disease-associated shifts and would mostly highlight study-specific differences. To examine whether MUSiCC can alleviate this problem, we again performed a principal coordinate analysis on the healthy samples from the three studies, but first using MUSiCC to correct the calculated abundances in the various samples. Indeed, in sharp contrast to the pattern observed in Figure 8A, we found that the study-based clustering was much less obvious (though still present) after the application of MUSiCC, with the healthy samples from the T2D and IBD studies largely overlapping and the distances between clusters becoming generally smaller than the distances within clusters (Figure 8B). Accordingly, we next examined whether MUSiCC could indeed help in the identification of disease-associated pathways when using the healthy samples from one study as controls for the cases from a different study. Specifically, we performed a comparative pathway analysis using the T2D cases together with either stool samples from HMP or the healthy samples from the IBD study as controls, and examined how many of the T2D-associated pathways identified with the original T2D dataset can be discovered (Methods). Overall, we found that using controls from a different study resulted in a much larger set of identified pathways, likely reflecting inter-study variation. We therefore restricted our cross-study analysis, only considering the same number of the most significant pathways detected with each dataset, to avoid artificially high recall levels (Methods). We found that when using compositional normalization, out of the 19 T2D-associated pathways identified with the original controls, only 2 (10.5%) pathways were recovered when using controls from HMP and only 1 (5.3%) pathway was recovered when using controls from the IBD study (Figure 8C). In contrast, when using MUSiCC, out of the 36 T2D-associated pathways detected with the original dataset, 8 (22.2%) pathways were recovered when using controls from HMP and 5 (13.8%) when using IBD controls (Figure 8C). Evidently, while inter-study variation is still a challenging problem, MUSiCC clearly and effectively improves our ability to pool samples from multiple studies and to increase the power to detect biological pathways associated with different diseases.

**Figure 8.**
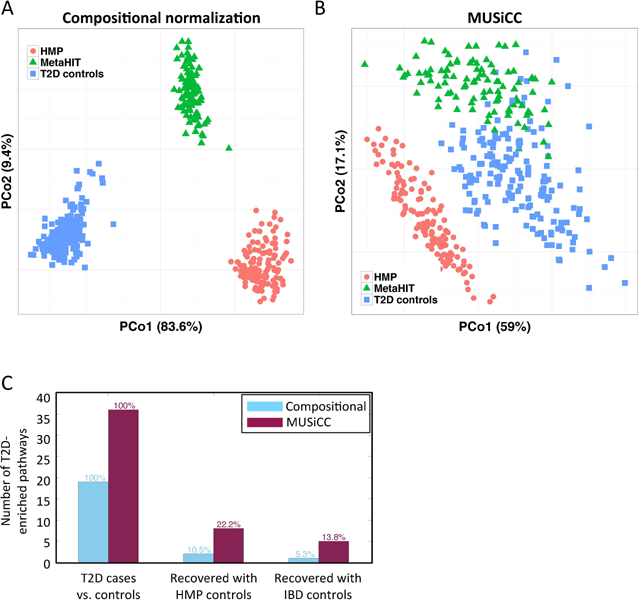
The impact of MUSiCC on pooling samples from multiple studies. **(A)** A principal Coordinate Analysis (PCoA) plot, illustrating the variation within the pathway-level functional profiles of healthy individuals from 3 independent studies of the human gut microbiome, using compositional normalization. Each dot represents a single sample and the proportion of variance explained by each of the first two principal coordinates is indicated on the axes’ labels. **(B)** A similar PCoA plot after the application of MUSiCC. **(C)** The number of T2D-associated pathways that are recovered (see Methods) when using as control the original T2D study control samples (left), HMP samples (middle), or IBD control samples (right). For each case, the number of pathways recovered with compositional normalization and with MUSiCC is illustrated.

### MUSiCC significantly reduces spurious variation in simulated bacterial samples

Above, we demonstrated that MUSiCC reduced spurious variation and improved our ability to detect disease-associated pathways in real metagenomic datasets. Such datasets provide a means to assess the full range of factors that potentially impact functional profile measurements. However, since the exact underlying taxonomic and functional compositions in these real datasets are unknown they cannot serve as a gold standard for evaluating our method or for comparison. We therefore wanted to further examine the ability of MUSiCC to reduce spurious variations on a synthetic dataset, where the true abundances of KOs and pathways are available. To this end, we generated 20 simulated metagenomic samples, each of which consisted of 500,000 reads generated at random from a mock community of reference genomes that were randomly assigned different relative abundances (see Methods and Table S5). We then mapped the reads in each sample to the KEGG database and calculated the read count of each KO in each sample. We finally normalized and corrected the obtained KO abundances using either MUSiCC or standard compositional normalization.

First, we compared the calculated abundance of each KO (using either MUSiCC or compositional normalization) with its real underlying average copy number in each sample, to examine the variation in estimated abundances across samples. We found that with compositional normalization, KOs with identical average copy numbers in different samples exhibit a wide range of normalized abundances across the 20 different simulated communities (Figure 9A). In sharp contrast, correcting KO abundances with MUSiCC resulted in a markedly narrower range of estimated abundance values for KOs with the same average copy number (Figure 9B). Moreover, as demonstrated in Figure 9B, MUSiCC not only reduced spurious inter-sample variation, but also provided an accurate estimation of the average copy number values of the various KOs, offering clear and biologically interpretable abundance values.

**Figure 9.**
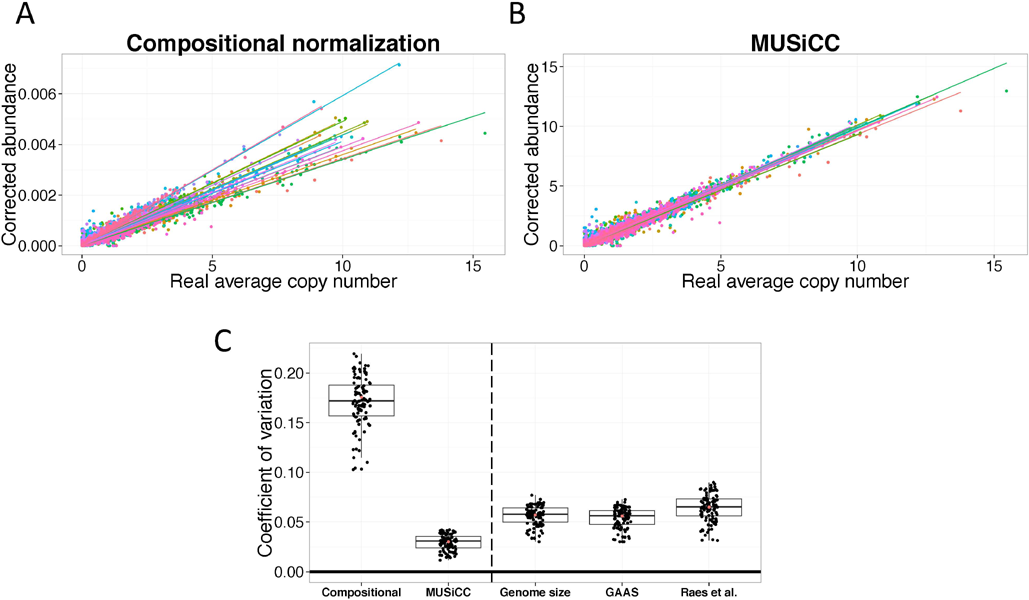
Evaluation of different normalization methods across 20 simulated metagenomic samples. Scatter plots are shown, comparing the average copy number of each KO in each sample to the corrected abundance of the KO in each sample as calculated by compositional normalization **(A)** and by MUSiCC **(B)**. Each sample is represented by a different color and a regression line for each sample is illustrated. The markedly reduced variability between the slopes of the regression lines in panel B (0.99±0.022; CoV=0.022 for MUSiCC vs. 2226±367; CoV=0.165 for compositional normalization) highlights the beneficial impact of MUSiCC in reducing spurious inter-sample variation and its overall accuracy in inferring underlying copy numbers. **(C)** Comparison of the observed coefficient of variation (CoV) in estimating the abundance of the flagellar assembly pathway across simulated samples using different normalization methods. Notably, in this set of samples, this specific pathway should exhibit no variation. Each dot represents the CoV calculated based on a random subset of 10 samples and the box plots illustrate the distribution of CoV values across 100 such subsets. Each box plot represents the 25^th^ and 75^th^ percentiles, with the median in black and whiskers extending to approximately ±2.7 standard deviations. The red dot represents the CoV calculated when using all 20 simulated samples.

Next, since many comparative analyses of metagenomes are performed at the pathway level (e.g., by summing the abundances of all the KOs associated with a given pathway), we wanted to specifically examine the impact of MUSiCC on spurious pathway abundance variation. To this end, in generating the simulated samples discussed above, we intentionally limited our selection of bacterial genomes to those that contain the entire set of flagellar assembly genes with the same exact copy numbers. Accordingly, any observed variation in the flagellar assembly pathway across the various samples is, by construction, spurious. Indeed (see Figure 9C), the coefficient of variation (*CoV;* standard deviation over mean) of the corrected abundance of this pathway across samples was markedly lower with MUSiCC (CoV=0.03) than with compositional normalization (CoV=0.175). To allow a more rigorous statistical analysis of the performances of each normalization method and to quantify the robustness of the various methods to subsampling, we further used a bootstrapping approach, repeatedly selecting a subset of the simulated samples and calculating the coefficient of variation observed in the abundance of this pathway across each subset. Comparing the distribution of CoV obtained for 100 subsets, we confirmed that inter-sample variation is indeed significantly lower with MUSiCC than with compositional normalization (Figure 9C; P<10^−72^, Student’s t-test for the difference in mean CoV). We additionally found that the distribution of CoV values obtained with MUSiCC across these 100 subsets exhibited lower variance than the distribution of CoV values obtained with compositional normalization (Figure 9C; P<10^−29^, F-test), suggesting that MUSiCC is also more robust to subsampling.

Finally, since our analysis revealed that variability in the average genome size across real samples is one of the factors contributing to spurious inter-sample variation, we wished to use the simulated samples described above to compare MUSiCC to normalization that is based on the average genome size in each sample. In addition to using the real average genome sizes (which are known for the simulated samples), we utilized two previously introduced methods, namely GAAS [22] and Raes *et a*l. [23], for metagenomic-based estimation of the average genome size in each sample (see Methods). While these methods were not developed with the aim of inter-sample normalization in mind, the average genome size estimates they provide may still be applied to correct observed spurious variation. We found that while both real and estimated genome size normalization markedly decreased spurious variation across samples in KO abundances (Figure S6) and in pathway abundances (Figure 9C), MUSiCC significantly outperformed these genome size based approaches. For example, the mean CoV for the abundance of the flagellar assembly pathway across samples was significantly lower in MUSiCC than in any genome size normalization methods (P<10^−50^, P<10^−50^, and P<10^−60^, for real average genome size, GAAS, and Raes *et al.* respectively; Student’s t-test), as was the variance in the CoV cross the 100 subsets described above (P<0.02, P<0.03, and P<10^−4^, for real average genome size, GAAS, and Raes *et al.* respectively; F-test). These results strengthen our finding that other sample-specific properties impact inter-sample variation and suggest that our USiCGs-based approach indeed captures a wide range of determinants in reducing spurious variation (and see also the discussion below), providing a more reliable characterization of the functional capacity of different metagenomes.

## Conclusions

The development of computational and statistical protocols for analyzing high-throughput metagenomic data has been a major line of research in recent years. Such protocols are essential for accurately characterizing the composition of various microbial communities and ultimately for reliably detecting functional shifts that may be associated with disease. These protocols often include a compositional normalization step, wherein read counts are converted into relative abundances, allowing researchers to compare different samples with potentially markedly different coverage. Importantly, this normalization procedure is extremely common and has become the *de-facto* standard in metagenomic assays and especially in human microbiome studies. Above, however, we systematically demonstrated that this normalization scheme introduces marked inter-sample spurious variation both across samples from the same study and across different studies of the human microbiome. We further rigorously characterized various sample-specific properties that give rise to this variation, demonstrating that the average genome size, the species richness, and the average genome mappability in the sample all independently contribute to spurious inter-sample variation. Rather than trying to reverse these complex and intertwined effects, we suggested a simple correction method, using a large set of universal single-copy genes to normalize the abundance of all other genes. This method accordingly not only corrects spurious variation across samples and facilitates accurate downstream comparative analyses, but also provides a more biologically relevant and meaningful abundance measure, estimating the average genomic copy number of every gene in the sample.

We additionally demonstrated that various gene-level properties, such as sequence conservation and nucleotide content, induce measurable and clear spurious intra-sample variation, further biasing calculated gene abundance profiles. We showed that the impact of these gene-specific properties on the measured abundance of universal single-copy genes can be modeled through an L_1_-regularized linear regression approach, and that the learned model can be used to partially correct such spurious intra-sample variation across all other genes in the metagenome. Notably, correcting intra-sample variability can be done by either using the built-in intra-MUSiCC trained model or by learning a specific model *de novo* for each new sample. Since our built-in model was learned using the HMP samples, this model is especially appropriate for samples that were obtained through a similar protocol. For samples obtained through different technologies or different processing pipelines that could potentially give rise to different biases, the intra-MUSiCC model should be ideally retrained on each dataset. Importantly, however, model training is extremely fast (see Methods), and since it is based only on the observed abundances of USiCGs (rather than on raw reads), running time is independent of the number of reads or their length.

Correcting inter-and intra-sample biases is clearly crucial for obtaining an accurate measure of gene abundance and for faithfully characterizing the functional profile of the metagenome. Since the true functional profile of the human microbiome is generally unknown, we used multiple approaches to confirm that our correction methods indeed yield improved abundance values and better recover the expected abundance and correlation structure of various sets of genes. Most importantly, however, we demonstrated that our methods not only provide a more accurate gene abundance measure, but also has a significant impact on the discovery of differentially abundant genes and of enriched pathways. Specifically, we showed that our MUSiCC pipeline markedly enhances the identification of disease-associated pathways, offering substantially increased statistical power both in terms of the number of pathways identified and the number of samples required for identification. Perhaps most remarkably, we found that MUSiCC facilitates efforts to pool data from several human microbiome studies, correcting much of the study-specific variation and laying the foundation for future cross-study comparative analyses. Notably, without MUSiCC, such study-specific variation was so dramatic that it masked practically any genuine, disease specific variation in the data.

In addition, we also evaluated MUSiCC using a set of simulated samples in which the underlying copy number of each gene in each sample is known. We demonstrated that MUSiCC significantly reduced spurious inter-sample variation both at the KO level and at the pathway level compared to compositional normalization. We further demonstrated that using the average genome size in each sample (either real or predicted) for normalization is not sufficient for removing inter-sample variation and that MUSiCC significantly outperformed such a normalization approach even in the ideal case of simulated communities. This finding highlights the benefit of a marker gene based normalization scheme and specifically the advantage of using USiGCs as a yardstick for copy number estimation, since it can capture the full range of factors that may introduce inter-sample variation. Moreover, it should be noted that methods for estimating average genome size often employ scenario-specific optimized parameters or require complete alignment, whereas MUSiCC is a parameter-free method that relies solely on KO measured abundances.

Clearly, the development of robust computational and statistical methods for an accurate characterization of gene abundances is an ongoing effort. Specifically, while inter-MUSiCC potentially corrects most of the inter-sample variation introduced by sample-specific properties, correcting gene-specific intra-sample biases is a much more challenging task. For one, the set of gene-specific properties that could affect the measured abundance of a gene may be markedly larger than the set we analyzed. Moreover, the directionality and magnitude of such effects may not be consistent across different groups of genes (note, for example, that a USiCGs-based model corrected >50% of the variation in unseen USiCGs, but only ~17% of the variation in semi-USiCGs; and see Ref. [25]). Distinguishing gene-specific effects from sampling noise is also challenging. We therefore hope that future work could further elucidate the contribution of additional properties and fine-tune the model relating gene-specific features to functional metagenomic profiling biases. Another opportunity for future extension of our method is going beyond bacterial and archaeal organisms and accounting also for other domains of life. Recent studies, for example, have highlighted the importance of the fungal component in the human microbiome [26]. Since our method was designed primarily with bacterial and archaeal genomes in mind, it may not be optimized to fungi-rich samples and may not accurately correct the abundances of fungal genes. Importantly, however, although the set of USiCGs used in our study was selected based on prokaryotic genomes, it is in fact well represented also in fungi, with 55 of the 76 USiCGs found on average in 58 of the 71 (82%) fungal genomes currently in KEGG, with a median copy number of 1.03. Rigorous processing of fungi-rich samples would thus ideally involve a modified and carefully selected set of marker genes and specifically tailored settings. Similarly, several recent studies have demonstrated that specific body sites, such as the skin, can harbor a substantial viral component [26–28]. Clearly, the human virome is an active and important component of the human microbiome, and as future efforts provide better characterization of this component, our methods could be further refined to account for the viral genes that may be identified across samples. One approach, for example, would be to augment our analysis with a sequence-based preprocessing step to distinguish reads that originated from this viral component. Combined, such extensions will allow researchers to improve various metagenomic processing pipelines and move towards a more accurate estimation of the functional composition of the metagenome. We believe that the MUSiCC framework and the improved estimation of the average genomic copy number of each gene in the metagenome represent an important step in this direction and can ultimately aid in the discovery of microbiome-based therapeutic targets.

## Methods

### Software implementation and distribution

MUSiCC was implemented in Python and is available for download or as a web-based application at: http://elbo.gs.washington.edu/software.html. In addition, it is available both in GitHub (https://github.com/omanor/MUSiCC) and as a pip-installable python package (i.e., *pip –U MUSiCC*). Input files for MUSiCC should be processed for host contamination removal and annotated by the KEGG orthology group database.

### Metagenomic data

Functional metagenomic data from multiple body sites were obtained from the Human Microbiome Project (HMP) [2] and were downloaded from the Data Analysis and Coordination Center (DACC) website: http://www.hmpdacc.org/. Gene relative abundance data were downloaded from ftp://public-ftp.hmpdacc.org/HMMRC/kegg_kos.pcl.gz. Genes were labeled by their KEGG Orthology (KO) Group and relative abundances were normalized for length. Sample IDs and body site information were downloaded from http://www.hmpdacc.org/HMASM/HMASM-690.csv. Sample and gene (KO) abundance data for the inflammatory bowel disease study [4] were obtained from a subsequent analysis of these samples performed by Ref. [29]. Sample and gene (KO) abundance data for the type 2 diabetes study were obtained from Ref. [11] and downloaded from http://gigadb.org/dataset/view/id/100036/files_page/21/samples_page/7.

### PICRUSt data

PiCRUSt [24] pre-calculated matrices for operational taxonomical units (OTUs) and their predicted KOs were downloaded from the developer’s github site: http://picrust.github.io/picrust/picrust_precalculated_files.html. OTU abundances mapped to GreenGenes IDs in the HMP body sites were obtained through personal communication with the lead author of PICRUSt.

### Detecting Universal Single-Copy Genes (USiCGs)

The list of USiCGs was compiled to include KOs that are both universal and appear in a single-copy in each genome. To determine the level of universality required, we used the list of 31 marker genes from the PhlyoSift pipeline [30], and examined the number of KEGG genomes in which each of these genes appear. We found that at minimum (after removing one outlier) these genes appear in 91.5% (2313) of the bacterial and archaeal genomes in KEGG, and therefore considered as universal any gene that appears in at least 91.5% of KEGG genomes. Of these genes, we further selected those that had an average number of copies per genome <1.1 in the genomes they appeared in, resulting in a list of 76 genes.

### Estimating average genome size in HMP samples

To estimate the average genome size in each HMP sample, we used PICRUSt pre-calculated files (see above), listing the predicted set of KOs and their copy number in each OTU. We multiplied the KO copy number by the average KO length in the KEGG database [1, 3] to obtain an estimate of the effective genome size of each OTU. We then computed the weighted average genome size in each sample, weighting the estimated genome size of each OTU by its relative abundance.

### Estimating the species richness and genome mappability in HMP samples

To estimate the species richness of a given sample, we counted the number of OTUs identified in that sample. To estimate the average mappability of short reads in a given sample, we utilized a recently introduced measure (implemented as part of the PICRUSt software [24]), termed *Nearest Sequenced Taxon Index (NSTI),* which aims to evaluate the evolutionary distance between each OTU in a sample to its closest reference genome. Specifically, we used PICRUSt pre-calculated NSTI values (see above) of each OTU, and for each sample computed the weighted average of (1-NSTI) across all OTUs in the sample, weighted by their relative abundance.

### Controlled correlations between USiCGs’ abundances and sample-specific properties

When calculating the correlation between the abundance of USiCGs and the various sample-specific properties, we used partial correlation analysis to control for the effect of other sample-specific properties. Partial correlations (and the corresponding P-values) were calculated using the “*partialcorr*” function in MATLAB. We additionally examined the correlation between species richness (and genome mappability) and USiCGs’ residuals after correcting for average genome size (see Figures S2C-D, S3E-F). Specifically, we used regression analysis with the median abundance of USiCGs in each sample as the response and the average genome size as the regressor, using the “*robustfit”* function in MATLAB to obtain the residuals. P-values for the correlation with the residuals are reported in Figure S2C-D and in Figure S3E-F.

### Identifying and analyzing OTU-Specific Genes (OSGs)

For each sample, we used the PICRUSt precalculated file (see above) that describes the predicted KO content of each OTU [24], combined with the list of OTUs that appear in the sample, to generate a list of KOs that are predicted to originate from only a single OTU in that sample (termed OTU-Specific Genes, or OSGs). For each OTU and each of its respective OSGs, we computed the Pearson correlation between the relative abundance of the OTU and the abundance of the OSG (using either compositional normalization or Inter-MUSiCC) across HMP stool samples. To be included in this analysis we required that both the OTU and the OSG were present in at least 5 samples. We then compared the correlation coefficients obtained when using compositional normalization to those obtained when using Inter-MUSiCC. To control for various factors that may confound the relationship between OSG and OTU abundances, we further performed a ratio-based analysis. For a pair of samples, sharing at least one OTU that contains OSGs, the fold-change between the OTU across the two samples was compared to the fold-change of each of its OSGs across the two samples. To avoid transitivity effects between pairs of samples, this process was performed as follows: for each OTU, the samples were ordered by the OTU’s relative abundance, and pairwise comparisons were performed only between consecutive samples in this ordered list. This was repeated for all OTUs. The distribution of the ratios between OSG and OTU fold-changes obtained when using compositional normalization was compared to the distribution obtained when Inter-MUSiCC was applied. The Statistical significance of the reduction in the mean and variance of this distribution were computed using the “*ttest*” and “*vartest2*” functions in MATLAB.

### Examining pairwise gene correlation structure with compositional normalization and with Inter-MUSiCC

We first downloaded all fully sequenced and annotated genomes from KEGG (downloaded on 07/15/2013). For each gene (*i.e.,* KO), we computed the Jaccard similarity between its presence/absence pattern across these genomes and the presence/absence pattern of all other genes. For each gene, we then also computed the Pearson correlation between its abundance across HMP stool samples and the abundances of all other genes. Given these two metrics, finally, for each gene, we computed the Pearson correlation between its correlation with all other genes across genomes and its correlation with all other genes in metagenomes. Distances and statistical significance were computed using the “*pdist*” and “*ttest*” functions in MATLAB.

### Gene-specific properties

Gene length and species statistics properties were downloaded from KEGG. Gene length was calculated as the average length of all genes labeled with the associated KO. Conservation and alignment properties were calculated by first downloading the gene sequences associated with each KO from KEGG. Next, MAFFT [31] was run on the set of genes for each KO, and statistics were calculated from the MAFFT multiple alignment output. Nucleotide content properties (e.g., mean GC%) were calculated from the set of gene sequences assigned to each KO in KEGG. KO recall and precision properties were obtained from a large-scale study of short read annotation (Carr and Borenstein, submitted) and describe the average recall and precision (across multiple genomes) in annotating simulated short shotgun reads originating from each KO.

### Regularized linear model linking gene-specific properties to intra-sample variation

For a given sample, we first defined the observed response as the fold-change between the abundance of each USiCGs and the mean abundance of all USiCGs in the sample. The model covariates were defined for all samples as the standardized values of the various USiCGs’ gene-specific properties (Table S2). When learning the model for a specific sample, we used a strict 5-fold cross-validation (CV) scheme to learn an Elastic-Net regularized linear model [32–34] that predicts the fold-change of each USiCG in the sample. Importantly, in each CV partition to training and test sets, we first learned the penalty parameter and model weights using solely the training data (by using an internal CV scheme with the MATLAB version of glmnet [32]), and evaluated the performance of our learned model by quantifying the fraction of variation the model explains when predicting the fold-change of USiCGs held-out as test data. Running time for the learning step on a typical sample (~13,000 KOs) took less than a second on a single core processor.

### Identifying Semi-Universal Single-Copy genes (semi-USiCGs)

Semi-USiCGs were selected as genes that were present in at least 2148 (85%) of bacterial and archaeal genomes in KEGG with an average number of copies per genome <1.1, but did not reach the USiCGs threshold of 2313 (91.5%) genomes, resulting in a list of 72 such genes.

### Identifying highly-correlated genes across genomes

For all KO pairs, we computed the Jaccard similarity across all KEGG genomes. Only the KO presence/absence pattern was considered, ignoring KO copy number. KO pairs with Jaccard similarity > 0.95 (*i.e.,* they agree on at least 95% of the genomes they appear in) were selected, resulting in 1074 such pairs. Clusters were defined as sets of 5 or more genes, all with pairwise Jaccard similarly > 0.95.

### Identifying differentially abundant genes

For a given gene and two sets of metagenomic samples (e.g., stool vs. tongue or IBD cases vs. controls), we compared the distribution of abundances between the two sets using the Wilcoxon rank-sum test. Genes that passed the Bonferroni correction (for comparing HMP body sites) or the FDR correction (for T2D and IBD cases vs. controls) with corrected p-values <0.05 were defined as differentially abundant.

### Identifying disease-associated pathways

For a given pathway and two sets of metagenomic samples (e.g., disease cases and controls), we first computed the pathway abundance in each sample as the sum of the abundances of genes associated with that pathway. Next, we computed an association p-value for the pathway by comparing the distribution of pathway abundance values in the two sets using the Wilcoxon rank-sum test. Pathways that had a higher median in disease samples and passed FDR correction (<0.05) were defined as disease-associated pathways.

### Evaluating an alternative normalization approach

To compare the ability of MUSiCC to discover disease-associated pathways with an alternative normalization approach previously used to process a set of oceanic samples [19], we followed the scheme applied in Ref. [19] to correct the gene abundances in the IBD dataset. Specifically, we first estimated the average genome size within each sample, using the set of 8 universal genes suggested in Ref. [19]. For each gene we estimated the average genome size, *G,* as *G=(g+L-2m)/f,* where *f* denotes the relative abundance of the gene in the sample, *L* denotes the sequence read length in the data (set to 75bp for the IBD dataset), and *m* denotes a minimum overlap parameter (set to 90 as in [19]). Next, we averaged the 8 estimations and corrected gene abundances by multiplying the relative abundance of each gene by this estimated average genome size of the sample. Finally, we used these corrected abundances and applied the same pathway-level comparative analysis to identify IBD-associated pathways. Since this normalization approach requires raw read counts, we could not apply it the other datasets analyzed in our study.

### Measuring T2D-associated pathway recovery when using T2D cases with HMP or IBD controls

First, we identified the original T2D-associated pathways as described above. Next, we repeated the same process but as controls used either HMP stool samples or healthy samples from the IBD study. To compute the number of recovered pathways, we counted the number of pathways that were discovered both in the original setting and in the cross-study setting. In order to prevent a situation where high recovery stems from the identification of many pathways, we limited the number of discovered pathways in the cross-study setting to be the number of pathways discovered in the original setting.

### Simulating microbial samples

To evaluate the performance of MUSiCC on a dataset in which the underlying KO and pathway abundances are known, we generated a dataset of synthetic metagenomic samples, following the procedure described in [35]. Specifically, we generated 20 simulated metagenomic samples, each of which consisted of 500,000 101bp reads generated at random from a collection of reference genomes that were randomly assigned different relative abundances (up to 100 fold) in each sample. To facilitate analysis of pathway level variation, we limited the set of genomes used to 21 bacterial genomes from KEGG that contained the entire set of KOs associated with the flagellar assembly pathway (Table S5), and each sample harbored 10 genomes randomly selected from this set. We then mapped the simulated reads to known KOs and calculated the read count of each KO in each sample, as previously described [35]. The underlying true average copy number for each KO was calculated based on the genomes included in each sample and weighted by their relative abundances.

### Using average genome size estimation methods for sample normalization

We utilized two previously introduced methods for estimating the average genome size in each simulated sample. First, to apply the method introduced in Raes *et al.* [23], we calculated the marker gene density for each sample. To allow the processing of these samples in a practical and reasonable time, we used our mBLASTx mapping to the KEGG database rather than re-blasting to the STRING database. As marker density, we used the total number of bases from all reads that matched any marker gene, divided by the length of the gene for each read mapped. Next, we plugged the resulting marker gene density, *x,* into the following equation for calculating the effective genome size (EGS):

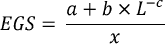

where *L* is the read length (101bp), and a, b, c are parameters optimized in the original study (*a*=21.2, *b*=4230, *c*=0.733) as described in Raes *et al.* [23]. The effective genome sizes obtained were highly correlated with the real average genome sizes in each sample (*r*=0.99; Pearson correlation). Using other sets of parameters described in the original study did not improve this correlation. Finally, we used these calculated effective genome sizes to normalize the measured KO abundances in each simulated sample.

Second, to apply the GAAS method introduced in [22], we first installed the GAAS software (version 0.17) from <u>http://sourceforge.net/projects/gaas/</u>. Since GAAS requires the complete BLAST alignment files, we ran BLAST for each simulated sample against the 21 genomes in our simulation (this represents both a best case scenario and is feasible computationally), using the same parameters as described in the original study: blastn -query *<sample fasta file>* -db *<simulation genomes nucleotide database>* -out *<GAAS blast results file>* -outfmt 6 -evalue 0.001 -gapopen 5 -gapextend 2 -word_size 11 -penalty -3 -reward 1. Next, we ran GAAS on the alignment results with the command: gaas -f *<sample fasta file>* -d *<GAAS nucleotide database>* -m *<GAAS blast results file>* -gt 0 -gp 0 -gs 0 -j 1, and obtained GAAS estimated average genome size in each sample.

Overall, the median error in average gnome size estimation was <1%, comparable to the accuracy reported in the original study. These GAAS average genome size estimations were again used to normalize the measured KO abundances.

## Acknowledgements

We thank Rogan Carr for valuable advice and for data on the mappability of genes used in this work. We thank Morgan Langille for information about the PICRUSt pipeline. We are grateful to all members of the Borenstein lab for helpful discussions. We thank the reviewers for their in-depth review and their suggestions for the improvement of our manuscript and software. This work was supported in part by NIH P30 DK089507 and by New Innovator Award DP2 AT007802-01 to EB.

## Figure Legends

**Figure S1.** Spurious inter-sample variation across HMP metagenomic samples in various body sites. See Figure 1B for definition of box and whisker plot.

**Figure S2.** Spurious variation in the abundance of USiCGs across HMP stool samples is correlated with **(A)** the species richness in the sample; R=0.44, P<10^−3^, and with **(B)** the average genome mappability; R=-0.85, P<10^−17^. **(C-D)** These correlations still hold after correcting the median USiCGs abundance and using the residuals with respect to the average genome size (R=0.35, P<0.005, and R=-0.33, P<0.008, for species richness and average genome mappability, respectively). Each point represents a single stool sample. Regression lines are illustrated in black. See Methods for more details on estimating average genome size, species richness, and genome mappability.

**Figure S3.** Spurious inter-sample variation in HMP tongue dorsum samples is correlated with sample-specific properties. **(A)** The relative abundance of USiCGs across HMP tongue dorsum samples. See Figure 1B for definition of box and whisker plot. The median abundance of USiCGs across tongue dorsum samples is correlated with (**B**) the average genome size; R=-0.6, P<10^−3^, **(C)** the species richness in the sample; R=0.43, P<10^−3^, and **(D)** the average genome mappability; R=-0.85, P<10^−17^. See Methods for more details on estimating sample-specific properties. **(E-F)** The correlations with species richness and genome mappability still hold after correcting the median USiCGs abundance with respect to the average genome size and using the residuals (R=0.33, P=0.06 and R=-0.67, P<10^−4^, respectively). Regression lines are illustrated in black.

**Figure S4.** The impact of MUSiCC on the discovery of differentially abundant genes between HMP stool samples and samples from other HMP body sites, including **(A)** supragingival plaque, **(B)** buccal mucosa, **(C)** anterior nares, **(D)** posterior fornix, and **(E)** retroauricular crease. Venn diagrams are defined as in Figure 6.

**Figure S5.** Comparing the impact of standard compositional normalization, an alternative normalization approach [19], and MUSiCC on the discovery of disease-associated pathways. Pathways identified to be associated with inflammatory bowel disease using any of these three methods are illustrated. Bars denote the significance level of the association. The dots to the right of each bar indicate whether this association reached significance with FDR<0.05 with compositional normalization (cyan), the alternative normalization approach [19] (green), or MUSiCC (maroon).

**Figure S6.** Evaluation of different average genome size based normalization methods across 20 simulated metagenomic samples. Scatter plots are as in Figure 9 using the real average genome size **(A)**, the average genome size estimated by GAAS **(B)**, or the average genome size estimated by Raes *et al.* **(C)**, for normalization.

